# CCMetagen: comprehensive and accurate identification of eukaryotes and prokaryotes in metagenomic data

**DOI:** 10.1101/641332

**Authors:** Vanessa R. Marcelino, Philip T.L.C. Clausen, Jan P. Buchmann, Michelle Wille, Jonathan R. Iredell, Wieland Meyer, Ole Lund, Tania C. Sorrell, Edward C. Holmes

## Abstract

High-throughput sequencing of DNA and RNA from environmental and host-associated samples (metagenomics and metatranscriptomics) is a powerful tool to assess which organisms are present in a sample. Taxonomic identification software usually align individual short sequence reads to a reference database, sometimes containing taxa with complete genomes only. This is a challenging task given that different species can share identical sequence regions and complete genome sequences are only available for a fraction of organisms. A recently developed approach to map sequence reads to reference databases involves weighing all high scoring read-mappings to the data base as a whole to produce better-informed alignments. We used this novel concept in read mapping to develop a highly accurate metagenomic classification pipeline named CCMetagen. Using simulated fungal and bacterial metagenomes, we demonstrate that CCMetagen substantially outperforms other commonly used metagenome classifiers, attaining a 3 – 1580 fold increase in precision and a 2 – 922 fold increase in F1 scores for species-level classifications when compared to Kraken2, Centrifuge and KrakenUniq. CCMetagen is sufficiently fast and memory efficient to use the entire NCBI nucleotide collection (nt) as reference, enabling the assessment of species with incomplete genome sequence data from all biological kingdoms. Our pipeline efficiently produced a comprehensive overview of the microbiome of two biological data sets, including both eukaryotes and prokaryotes. CCMetagen is user-friendly and the results can be easily integrated into microbial community analysis software for streamlined and automated microbiome studies.

## Introduction

Microbial communities in natural and host-associated environments commonly harbour a mix of bacteria, archaea, viruses and microbial eukaryotes. Bacterial diversity has been extensively studied with high-throughput sequencing (HTS) targeting 16S rDNA markers (Caporaso et al. 2011; Taberlet et al. 2012). However, these do not amplify eukaryotic sequences, and our knowledge on the diversity and distribution of microbial eukaryotes is limited (Bik et al. 2012; Norman et al. 2014). Although there is an increasing number of studies using eukaryotic-specific markers, these are relatively uncommon and face multiple methodological limitations (Piganeau et al. 2011; Marcelino and Verbruggen 2016). The problematic amplification step can be bypassed by sequencing the total DNA (metagenome) or RNA (metatranscriptome) in a sample to characterize all the genes contained or expressed within it. Metagenomics and metatranscriptomics are promising tools to bridge the knowledge gap in the diversity of microbial eukaryotes because they are essentially kingdom-agnostic, are less susceptible to amplification bias, and yield a large set of genes that can be used for taxonomic identification.

Multiple software packages have been developed to reveal the species composition of metagenomic samples (reviewed in Breitwieser et al. 2017). While well-known bacterial species can be easily identified at the species and strain levels (Truong et al. 2015; Scholz et al. 2016), it remains challenging to obtain a fine-grained taxonomic classification of lesser-known species and microbial eukaryotes (Sczyrba et al. 2017; Nilsson et al. 2019). Many of the current metagenomic classifiers assign a taxonomy to each short sequence read individually (Breitwieser et al. 2017). However, as closely-related species share very similar or identical genome portions, short reads often map to multiple species in the reference data set. Some metagenomic classifiers, like MEGAN (Huson et al. 2007) and Kraken (Wood and Salzberg 2016), address this issue by calculating the lowest common ancestor (LCA) among all species sharing those sequences. Paradoxically, this classification strategy is negatively affected by the increasing size of reference databases: as identical regions in reference databases become more common, fewer reads can be classified at the species level (Nasko et al. 2018). Other classifiers use a database of clade-specific diagnostic regions (e.g. Truong et al. 2015). While highly accurate, this procedure relies heavily on reference databases of complete genomes, which often cannot be readily updated by the end-user. Complete genomes are available for only a small fraction of the microbial eukaryotic species. For example, as of April 2019, the widely used NCBI RefSeq database contained 285 fungal genome sequences, even though it is estimated that there are over 2 million species of fungi (Hawksworth and Lucking 2017). Therefore, relying on these databases of complete genomes restricts the inclusion of microbial eukaryotes in metagenome studies.

A recently-developed concept in read mapping – the ConClave sorting scheme, implemented in the KMA software (Clausen et al. 2018) – is more accurate than other mapping strategies as it takes advantage of the information from all reads in the data set (Figure 1). Our goal was to use this approach to produce an accurate metagenomic classification pipeline that will allow the inclusion of microbial eukaryotes in metagenomic studies. We present a novel tool - CCMetagen (ConClave-based Metagenomics) – to process KMA sequence alignments and produce highly accurate taxonomic classifications from metagenomic data. We benchmark CCMetagen using simulated fungal and bacterial metagenomes and metatranscriptomes. Additionally, we use two case-studies with real biological data to demonstrate that CCMetagen effectively produces a comprehensive overview of the eukaryotic and prokaryotic members of microbial communities.

**Figure 1.**
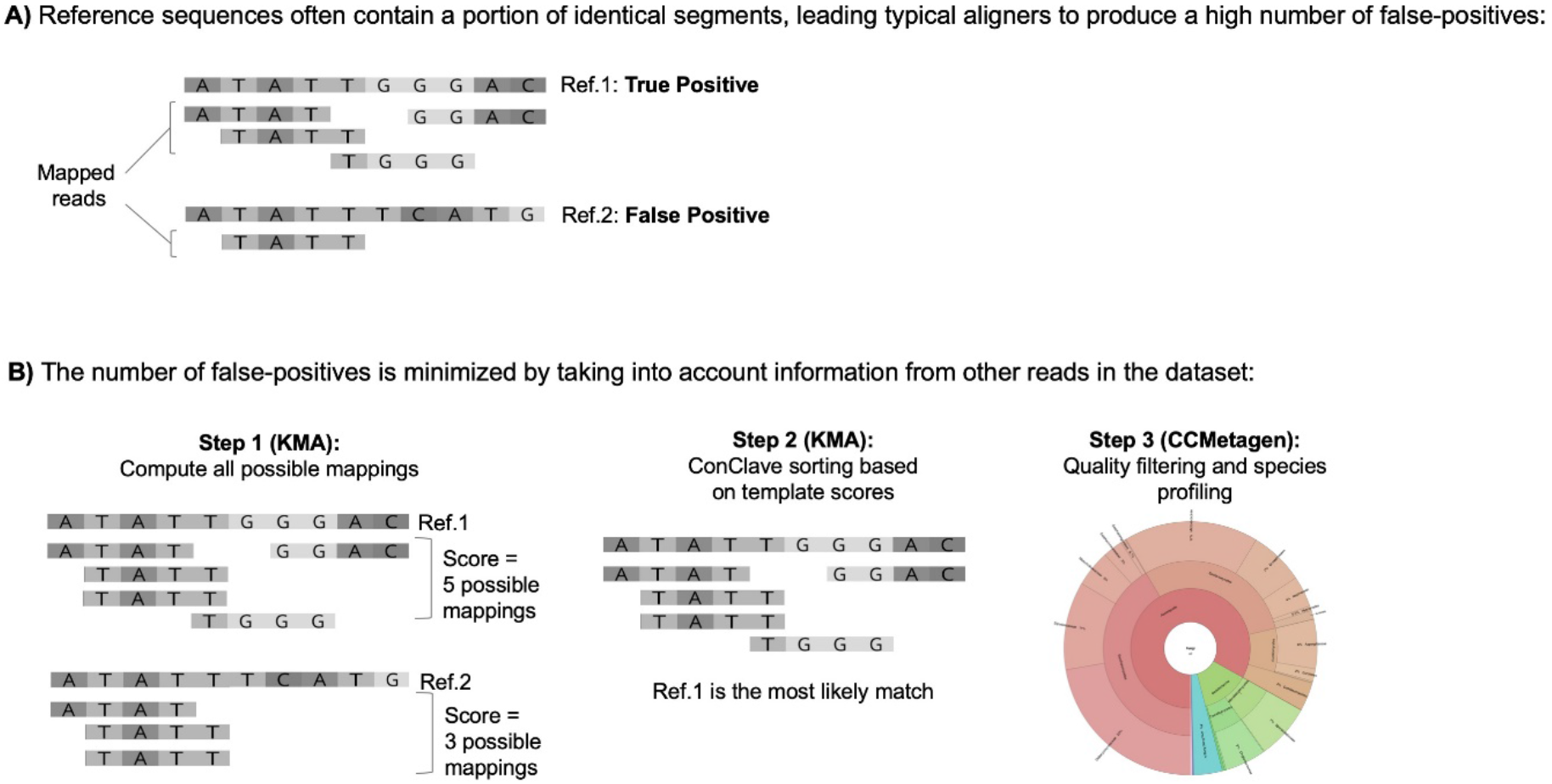
Overview of the ConClave sorting scheme applied to species identification in metagenomic data sets. The figure represents a data set containing 5 sequence reads (4bp) and two closely-related reference sequences (templates), including a true positive (Ref.1) and a potential false positive (Ref.2). (A) Commonly used read mappers yield a high number of false-positives because reads can be randomly assigned to closely-related reference sequences sharing identical fragments spanning the whole sequence read (represented by the ATATT region). (B) The KMA aligner minimizes this problem by scoring reference sequences based on all possible mappings of all reads, and then choosing the templates with the highest scores. Coupled with KMA, CCMetagen produces highly accurate taxonomic assignments of reads in metagenomic data sets in user-friendly formats.

## Results

### Implementation and availability

Metagenomic reads (or contigs) are first mapped against a reference database with KMA (Clausen et al. 2018), which implements the ConClave sorting scheme for better-informed and highly accurate alignments (Figure 1). CCMetagen is then used to perform quality filtering and produce taxonomic classifications that can be explored in text or interactive visualization formats (Krona plots - Ondov et al. 2011). Our pipeline uses the NCBI taxonomic database (taxids) to produce ranked and updated taxonomic classifications, so that the ever-changing species nomenclature issue is minimized (Federhen 2012). CCMetagen yields classifications at a taxonomic level that reflect the similarity between query and reference sequences. This ranked classification means that the method is able to identify species that have only distant relatives in reference databases (*e.g.* undescribed genera), as well as well-known microorganisms. The output of CCMetagen can be easily converted into a PhyloSeq object for statistical analyses in R (McMurdie and Holmes 2013). The pipeline is sufficiently fast to use the entire NCBI nucleotide collection (nt) as a reference database, thereby enabling the inclusion of microbial eukaryotes – in addition to bacteria, viruses and archaea – in metagenome surveys. Our program is implemented in Python 3 and is freely available at https://github.com/vrmarcelino/CCMetagen.

### Fungal classifications are more accurate with the CCMetagen pipeline

To test the performance of CCMetagen in identifying an important and diverse group of microbial eukaryotes, we simulated *in silico* a fungal metatranscriptome (15 species) and a fungal metagenome (30 species). We then benchmarked CCMetagen’s performance by comparing it with widely used metagenomic classification software, including Centrifuge (Kim et al. 2016), Kraken2 (Wood and Salzberg 2016) and KrakenUniq (Breitwieser et al. 2018). These programs were chosen because they are compatible with custom-made reference databases, which is a desirable flexibility when working with microbial eukaryotes. KrakenUniq was recently shown to outperform eleven other classification methods when using the NCBI nucleotide collection (‘nt’ database), including Diamond/Blast + MEGAN (Altschul et al. 1990; Huson et al. 2007; Buchfink et al. 2015), CLARK (Ounit et al. 2015), GOTTCHA (Freitas et al. 2015), PhyloSift (Darling et al. 2014) and MetaPhlAn2 (Truong et al. 2015). KrakenUniq therefore provides a gold standard for the available tools. We evaluated precision, recall and F1 scores of the benchmarked software in identifying fungal taxa in the simulated fungal metagenome and metatranscriptome (see Methods). The F1 score is the harmonic average of precision and recall; high F1 scores can be interpreted as a good trade-off between precision and recall.

The CCMetagen pipeline achieved the highest precision and F1 scores of all the approaches tested (Figure 2, Supplemental Table S1, Supplemental Figures S1 and S2). KrakenUniq achieved higher precision than Kraken2 and Centrifuge when using an ideal database (*i.e.* RefSeq-bf, which contains only the complete and curated genomes of fungi and bacteria, containing all species from the test data set). However, the performance of KrakenUniq decreased substantially when the database was incomplete (*i.e.* RefSeq-f-partial, where a part of the reference sequences was removed to mimic the effects of handling species without reference genomes).

**Figure 2.**
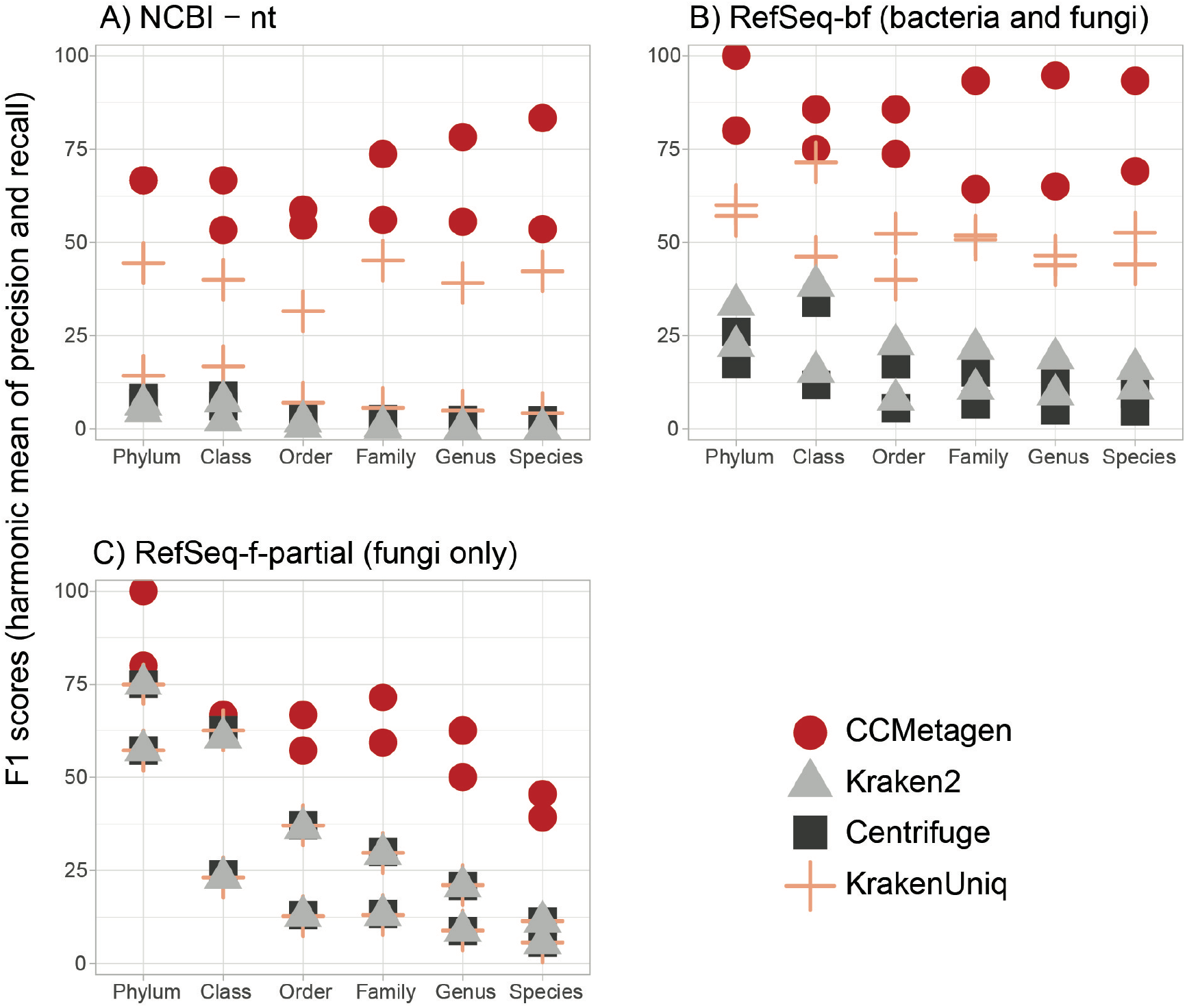
The CCMetagen pipeline has a higher F1 score than other metagenomic classification methods for all taxonomic ranks. The two points for each program and taxonomic rank represent the results using a simulated metagenome and a metatranscriptome sample of a fungal community. (A) Results using the whole NCBI nt collection as a reference database. (B) Results using the RefSeq-bf (bacteria and fungi) database, containing all bacterial and fungal genomes available. (C) Partial RefSeq database containing only some of the fungal species currently present in the RefSeq-bf database, mimicking the effects of dealing with species without representatives in reference data sets. In this case, Kraken2, Centrifuge and KrakenUniq have overlapping results. Refer to Supplemental Table S1 and Supplemental Figures S1 and S2 for more information, including precision and recall.

Centrifuge, Kraken2 and KrakenUniq yielded many more taxa than included in the test data sets: for example, Centrifuge, when used with the nt database, reported 6950 species in the simulated metagenome containing 30 species, while CCMetagen yielded only 15. Naturally, their recall was very high – Centrifuge and KrakenUniq recovered 100% of the taxa present in the test data set when using the RefSeq-bf and nt reference databases (Supplemental figure S2). The species-level recall of Kraken2 decreased when using the nt database. CCMetagen recovered between 50% and 100% of the species when used with RefSeq-bf and nt databases (Supplemental Table S1).

The fastest processing time was achieved by Kraken2 (Table 1). The combined CPU time of KMA and CCMetagen (*i.e.* the CCMetagen pipeline) was faster than Centrifuge and KrakenUniq when using the whole NCBI nt database, but it was the slowest approach when using the RefSeq database. The KMA indexing of the nt database was limited to only include *k-*mers with a two-letter prefix, which on average corresponds to only saving non-overlapping *k-*mers. This prefixing substantially increases the speed and could also be applied to the RefSeq database if faster processing time is required (Supplemental Materials). When the NCBI nt data set was used, CCMetagen required ~ 15min to process a sample (~5GB, 7.8M reads on average).

**Table 1.** CPU time (in minutes) required to analyze a simulated fungal metatranscriptome (mtt) and a fungal metagenome (mtg).

|  | nt |  | RefSeq-bf |  | RefSeq-f-Partial |  |
| --- | --- | --- | --- | --- | --- | --- |
|  | mtt | mtg | mtt | mtg | mtt | mtg |
| **Kraken2** | 10.92 | 7.05 | 5.29 | 3.98 | 4.48 | 3.50 |
| **CCMetagen\*** | 17.24 | 13.54 | 85.74 | 67.00 | 69.29 | 20.58 |
| **Centrifuge** | 40.11 | 27.54 | 23.70 | 19.41 | 16.67 | 16.10 |
| **KrakenUniq** | 74.11 | 74.94 | 43.33 | 40.85 | 29.65 | 21.04 |

\* The CCMetagen time was calculated as the sum of the CPU time used by KMA and CCMetagen.

### Bacterial communities are best depicted with the CCMetagen pipeline

We assessed the performance of the CCMetagen pipeline with 10 bacterial communities simulated at different levels of complexity (Segata et al. 2012; McIntyre et al. 2017). Using the NCBI nt collection as reference, CCMetagen achieved the highest precision and F1 scores, at all taxonomic ranks (Figure 3). Recall was highest for Centrifuge and KrakenUniq. In this data set, the recall of Kraken2 was higher than CCMetagen from phylum to family-level classifications, but lower than CCMetagen at genus and species level.

**Figure 3.**
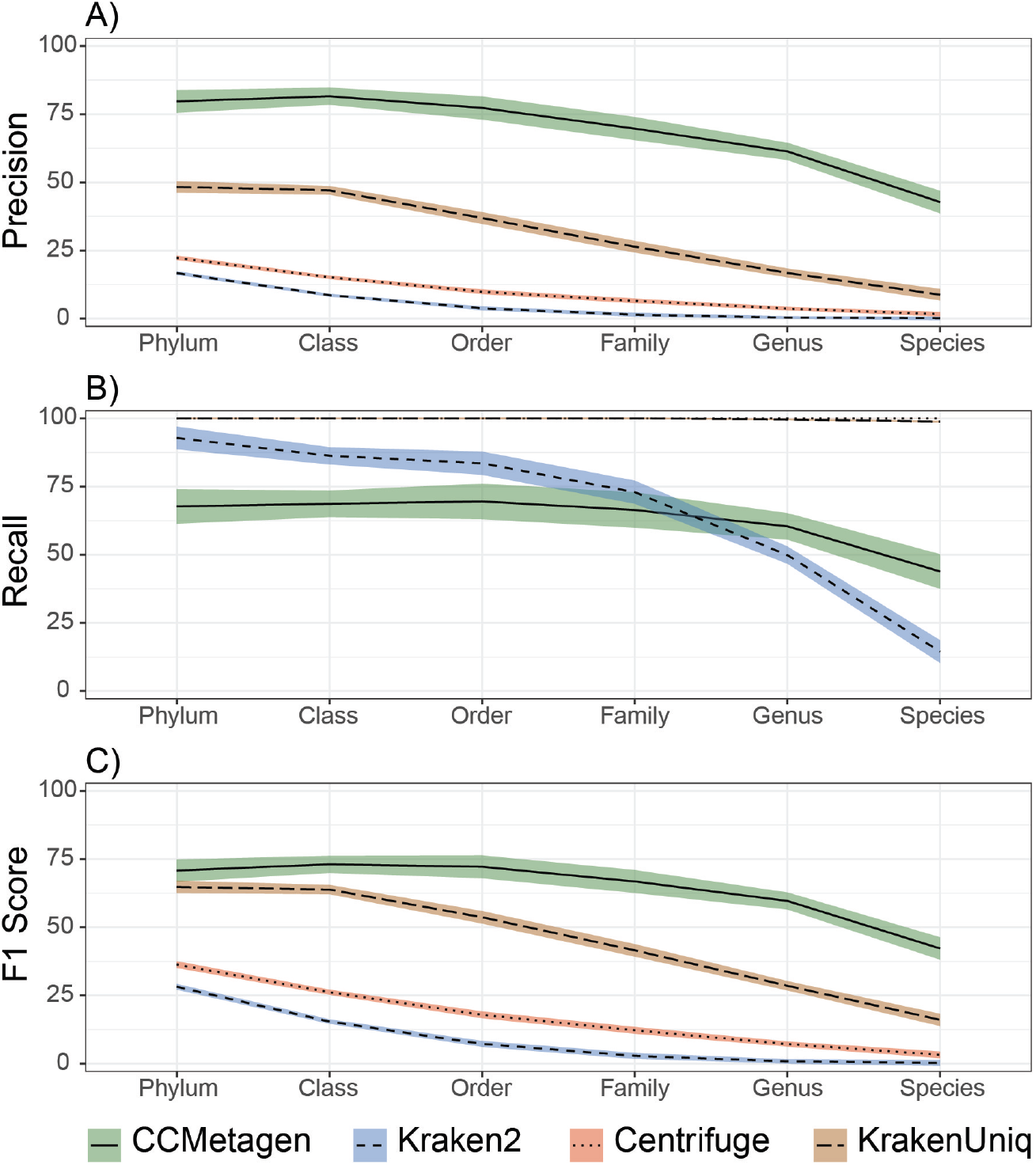
CCMetagen pipeline performance for bacterial classifications, compared with Kraken2, Centrifuge and KrakenUniq. Precision (% of true positives), Recall (% of taxa identified) and F1 scores represent averages across 10 simulated metagenome samples. Shaded areas indicate 75% confidence intervals.

The complete CCMetagen pipeline (KMA + CCMetagen) required an average of 2.1 minutes to process the bacterial metagenomes (+/− 0.26 SD). It was slower than Kraken2 (average 0.27m, +/− 0.21 SD) and faster than KrakenUniq (average 2.56m, +/− 2.60 SD) and Centrifuge (average 9.19m, +/− 0.80 SD).

### Biological data set 1: experimentally seeded fungal metatranscriptome

We validated the CCMetagen pipeline with a fungal community previously generated *in vitro* by culturing, processing and sequencing 15 fungal species (Marcelino et al. 2019a, Supplemental Table S2). The analyses were performed using the NCBI nt collection as reference. Our pipeline correctly retrieved 13 out of the 15 fungal species sequenced, in addition to identifying a small component of other eukaryotic (0.4%) and bacterial (3%) RNA, which likely represents laboratory contaminants (Figure 4, Supplemental Table S3).

**Figure 4.**
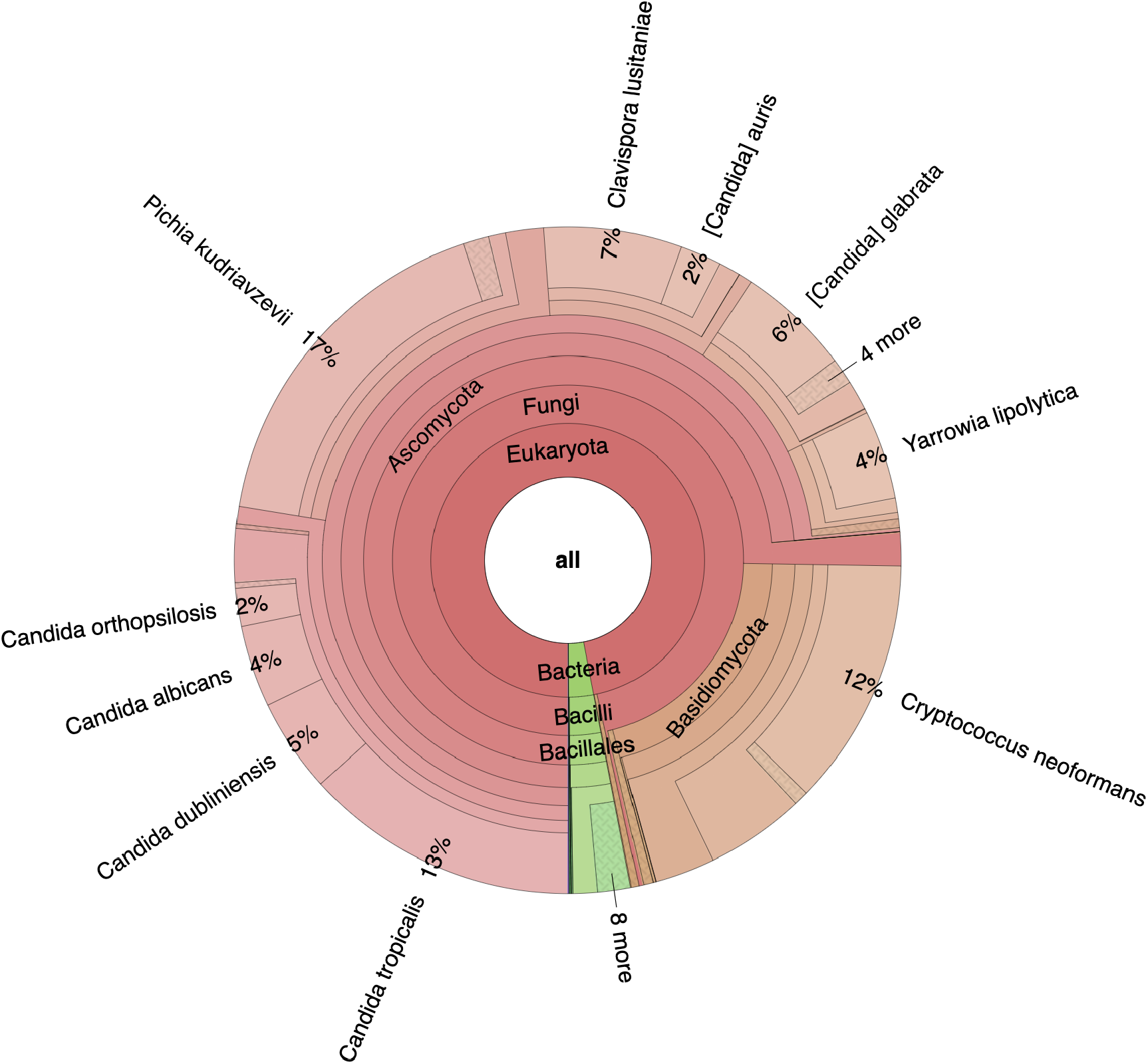
Snapshot of CCMetagen results for a spiked fungal community. This Krona graph shows the relative abundance of taxa at various taxonomic levels, which are color-coded according to their taxonomic classification at lower-ranks – here we see fungal taxa in shades of red, and bacterial taxa in shades of green. The Krona html file can be opened and interactively inspected in a web browser. Each circle represents a taxonomic level, where the user can click for a representation of the relative abundance at a given taxonomic rank. For a detailed list of taxa, refer to Supplemental Table S3.

As this data set contains the same 15 fungal species as those simulated *in silico*, it is possible to tease apart classification errors from laboratory-related confounders such as contamination. Accordingly, we were able to retrieve all 15 species when using the *in silico* data set, suggesting that the two false-negatives (*Schizosaccharomyces pombe* and *Debaryomyces hansenii*) were missing due to laboratory-related issues, such as RNA extraction biases, gene [under]expression and imprecise cell counts. We also identified seven times more false-positives in the seeded fungal metatranscriptome (44 species, while the simulated data yielded only 6). These additional 38 species were present at low abundance and most likely represent reagent and laboratory contaminants (Salter et al. 2014; Strong et al. 2014).

### Biological data set 2: the microbiome of Australian birds

We used the CCMetagen pipeline to characterize the gut microbiome associated with 9 metatranscriptome libraries from wild birds sampled at various sites across Australia (Wille et al. 2018; Marcelino et al. 2019b). Fungal and bacterial transcripts were observed in all libraries (Supplemental Table S4). Eukaryotic microbes accounted for 60% of the family-level diversity of the bird microbiome samples (taxa unclassified at family-level were not taken into account). Notably, fungi represented 12 of the 20 most abundant microbial families, surpassing the diversity of bacterial families (Figure 5). Among the fungal transcripts with a species-level classification, those attributed to the basidiomycete *Cystofilobasidium macerans* (Tremellomycetes) were the most abundant and were present in all bird libraries. Transcripts from species of *Mucor*, *Cladosporium*, *Metschnikowia, Fusarium* and *Cryptococcus* were common. Other microbial eukaryotes were also observed, including the trichomonad *Simplicimonas* and the Apicomplexan *Eimeria*. Archaeal and viral transcripts were also detected. The methanogenic archaea *Methanobrevibacter woesei*, which was previously reported in chicken guts (Saengkerdsub et al. 2007), was observed in two duck libraries. Influenza A virus was detected and confirmed with PCR-based methods (Wille et al. 2018). The CCMetagen results were parsed with PhyloSeq for a graphical representation of the most abundant microbes, and the R script to reproduce Figure 5 is available on the CCMetagen website.

**Figure 5.**
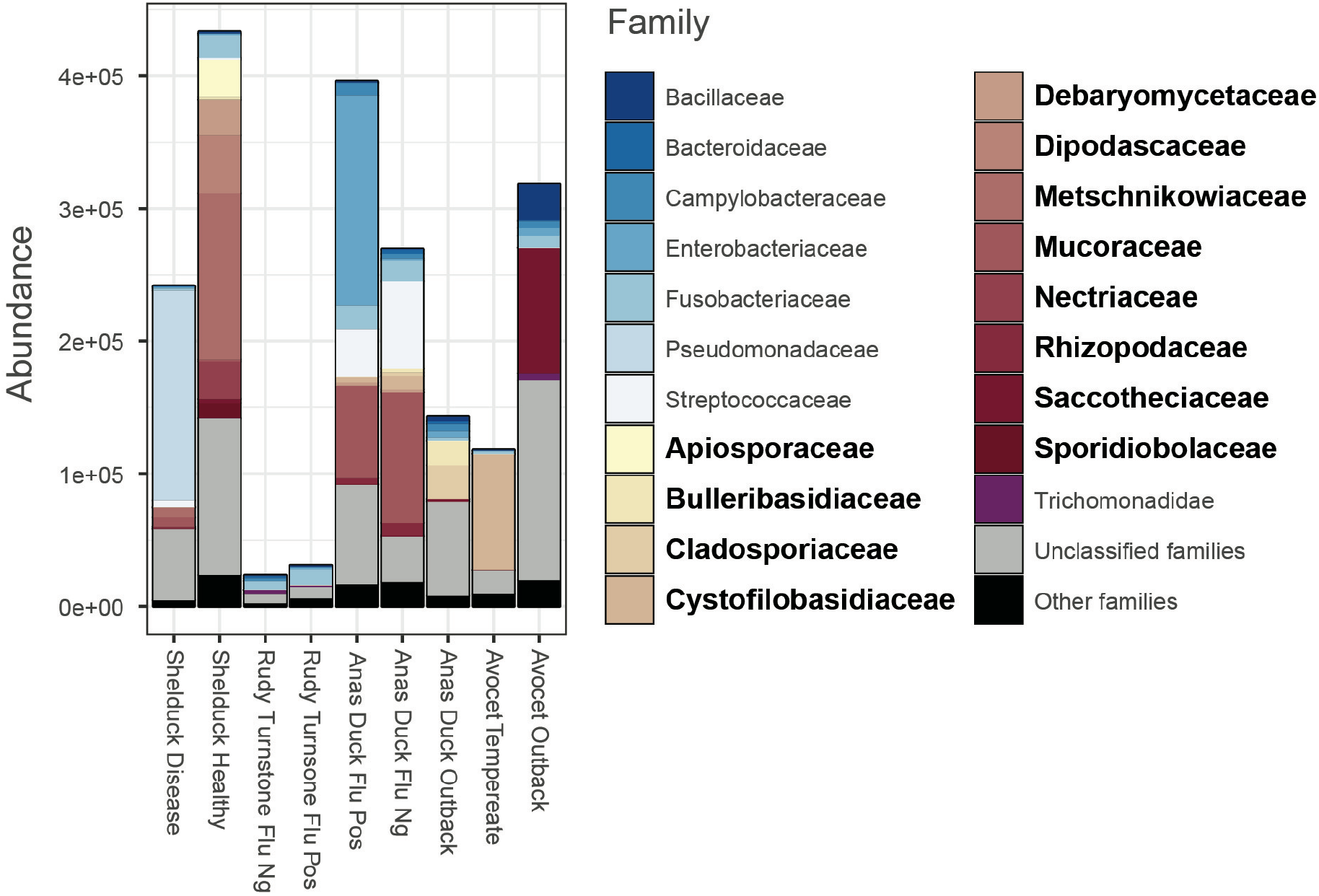
Microbial families in the microbiome of wild birds. The 20 most abundant families are shown, with fungal families indicated in bold. For a full list of taxa, refer to Supplemental Table S4. A tutorial and R scripts to reproduce these analyses are available on the CCMetagen website.

## Discussion

The application of the ConClave sorting scheme to differentiate highly similar genetic sequences (Clausen et al. 2018) represents an important step forward in metagenomic species profiling. We have applied this concept to develop a metagenome classification pipeline that is highly accurate yet fast enough to use the entire NCBI nucleotide collection as reference, thereby facilitating the identification of microbial eukaryotes in metagenomic studies. The species-level identifications of bacteria and fungi obtained with the CCMetagen pipeline were from 3× to 1580× more precise than other metagenome classifiers (across all databases tested). CCMetagen is therefore a powerful tool for achieving accurate taxa identifications across a range of biological kingdoms in metagenome or metatranscriptome samples.

Scarce reference data pose a major challenge to study any microbial system that is less well-studied than the human gut. Some of the methods with reportedly high accuracy rely heavily on reference databases of complete or near complete genomes. KrakenUniq, for example, showed relatively high precision and recall when using the RefSeq-bf database, which contained the complete genomes of all species in the test data set. However, when KrakenUniq was tested with an incomplete reference database (RefSeq-f-partial), the number of false positives increased, on average, from 51 to 221 species. This likely happens because it is relatively easy to identify a species that is present in the reference database, while it can be challenging to identify the closest match in the absence of a perfectly matching reference sequence. In the latter case, when reads are classified individually, multiple reference sequences can have identical levels of similarity, leading to a high number of false-positives. This is an obvious problem when working with microbial eukaryotes, for which very few complete genomes are available.

One of the many advantages of metagenomics is that it enables the detection of novel and rare microbes. Being able to distinguish between known and novel microorganisms in metagenomic data sets is a desirable feature possessed by surprisingly few metagenome classifiers. Some of these classifiers (e.g. MEGAN and Kraken) use the lowest common ancestor between all reference sequences that match the query sequence. The accuracy of these taxonomic classifiers tends to decrease as reference databases get populated with closely-related taxa (Nasko et al. 2018) and, paradoxically, well-known taxa can be classified at higher taxonomic ranks than rare or novel ones. CCMetagen classifies taxa at the lowest common ancestor that reflects the genetic similarity between the query and the reference sequence. As rates of molecular evolution can vary substantially among genes and species, it is currently not feasible to set a universal sequence similarity threshold that works equally well for all organisms and genes. By default, CCMetagen uses similarity thresholds previously determined for fungi (Vu et al. 2016; Vu et al. 2019). Importantly, CCMetagen allows the user to easily set different similarity thresholds or disable the threshold-filtering step entirely. While this strategy also has limitations, it is a better alternative to the reference-dependent method of calculating LCAs, even when using the default thresholds for bacterial classifications (Figure 3).

With CCMetagen, it is possible to confidently use metagenomics to identify microbial eukaryotes and prokaryotes in microbial communities. Our analyses of the gut microbiome of wild birds revealed an abundant and diverse community of micro-eukaryotes, representing 60% of the family-level diversity in the samples. We detected various species of *Mucor* and of basidiomycetes, including species of the opportunistic pathogen genus *Cryptococcus*. These and other non-ascomycetes fungi can be affected by mismatches in commonly used metabarcoding primers (Bellemain et al. 2010; Ihrmark et al. 2012; Tedersoo and Lindahl 2016). The fact that they were observed in high abundance indicates that metagenomics and metatranscriptomics are valuable for detecting these organisms in environmental samples. Importantly, CCMetagen can generate results in a format that resembles an Operational Taxonomic Unit (OTU) table that can be imported into software designed for microbial community analyses, such as PhyloSeq (McMurdie and Holmes 2013), facilitating downstream ecological and statistical analyses of the microbiome.

In summary, CCMetagen is a versatile pipeline implementing the ConClave sorting scheme (via KMA) to achieve highly accurate taxonomic classifications. The pipeline is fast enough to use the entire NCBI nt collection as reference, facilitating the inclusion of understudied organisms, such as microbial eukaryotes, in metagenome surveys. CCMetagen then produces ranked taxonomic results in user-friendly formats that are ready for publication (with Krona) or for downstream statistical analyses (with PhyloSeq). We expect that a range of novel ecological and evolutionary insights will be obtained as information about microbial eukaryotes in metagenomic studies becomes more accessible.

## Methods

### Test data sets

A fungal metagenome and a metatranscriptome were simulated *in silico* to assess the performance of CCMetagen and other classification pipelines in identifying the fungal members of a microbial community (Supplemental Table S2). Simulations were based on complete fungal genomes obtained from the NCBI RefSeq collection (Pruitt et al. 2007). The metagenome contained 30 fungal species and was simulated with Grinder (Angly et al. 2012) using parameters to mimic the insert size and sequencing errors of an Illumina library (-md poly4 3e-3 3.3e-8 -insert_dist 500 normal 50 -fq 1 -ql 30 10). Coverage was set to vary between 0.001× and 10× for different species. The metatranscriptome contained 15 fungal species and was simulated for a subsample of 4000 genes (CDSs) from each fungal genome. Transcripts were simulated with Polyester (Frazee et al. 2015), using the Illumina5 error model and gene expression following a normal distribution of average 3× (20% of genes up- and 20% down-regulated).

Additionally, 10 bacterial metagenomes simulated by Segata et al. (2012), and compiled in McIntyre et al. (2017), were used to assess the performance of the different classifiers in identifying prokaryotic communities with various levels of complexity. Each metagenome contained between 25 and 100 bacterial species.

### Reference databases

We used three reference databases: (i) “nt” - the NCBI nucleotide collection; (ii) “RefSeq-bf”, containing curated genomes of fungi (all assembly levels) and bacteria (only complete) in NCBI Reference Sequence Database; and (iii) “RefSeq-f-partial”, which is a subset of RefSeq-bf, containing only part of the fungal species in our test data sets. The RefSeq-f-partial database was built to assess how the programs perform when reference databases are incomplete, for example, when dealing with species without reference genomes. Fifteen species were removed, resulting in a database that contained 15 of the 30 species in the fungal metagenome sample, and 7 of the 15 species in the metatranscriptome sample (species removed from this data set are detailed in Supplemental Table S5). Details about databases download and indexing can be found in the Supplemental Material. The nt and RefSeq-bf databases indexed to function with KMA and CCMetagen are hosted in two sites, at https://cloudstor.aarnet.edu.au/plus/s/Mp8gLimDYoLfelH (Australia) and http://www.cbs.dtu.dk/public/CGE/databases/CCMetagen/ (Denmark).

### Benchmarking

Details about the quality control and data analyses are described in the Supplemental Materials. Metagenome classifications using Kraken2 v.2.0.6-beta, KrakenUniq v.0.5.6 and Centrifuge v.1.0.3-beta were performed using default values. The performance of the classifiers was assessed in terms of precision, recall, F1 score and CPU time. Precision was calculated with the formula:

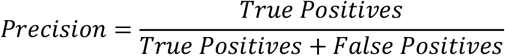

Recall was calculated with the formula:

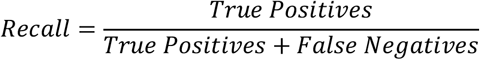

And the F1 score, which is the harmonic average of the precision and recall, was calculated as:

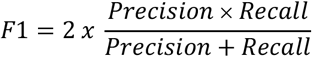

Precision and Recall were multiplied by 100 to indicate percentages. CPU time was calculated with GNU’s time function (user + sys). True and false positives, at several taxonomic levels (superkingdom to species), were calculated based on NCBI taxids. Only matches to organisms with valid taxids were included in the analyses. Valid but obsolete taxids (those that have changed due to nomenclature changes) were updated accordingly using the ETE toolkit (Huerta-Cepas et al. 2016). This strategy also minimizes nomenclature problems. For example: *Filobasidiella neoformans* is a life stage of *Cryptococcus neoformans*, they share a unique taxid (5207) regardless of the name attributed to the sequence in the reference database. The benchmarking scripts are available at: https://github.com/vrmarcelino/CCMetagen/tree/master/BenchmarkingTools.

### CCMetagen applied to real data sets

We validated the CCMetagen pipeline using two biological data sets: one defined fungal community (biological data set 1) and one set of environmental samples (biological data set 2). The fungal community was constructed by culturing, pooling and sequencing the same 15 fungal species used in the metatranscriptome simulated *in silico* (SRA BioProject number PRJNA521097) (Marcelino et al. 2019a).

The biological data set 2 consisted of nine metatranscriptome libraries derived from gut samples from Australian wild birds (SRA BioProject number PRJNA472212) (Wille et al. 2018). Quality control was performed as described in Marcelino et al. (2019b).

These samples were mapped to the NCBI nucleotide database using KMA with the options - 1t1 -mem_mode -and -apm f, and then processed with CCMetagen using default values. The results were parsed with PhyloSeq to produce a graph with relative abundances (Figure 5). A tutorial explaining the full analyses of the bird microbiome, from quality control to graphical representation with PhyloSeq, is available at https://github.com/vrmarcelino/CCMetagen/tree/master/tutorial.

### Data access

CCMetagen is freely available from https://github.com/vrmarcelino/CCMetagen (licensed under GNU General Public License v3.0). The simulated fungal metagenome and metatranscriptome sequence is available at https://cloudstor.aarnet.edu.au/plus/s/Mp8gLimDYoLfelH

## Acknowledgments

We thank the High Performance Computing support and the Research Data Support at the University of Sydney, and the Genomic Facilities at the Westmead Institute for Medical Research. We thank Mick Watson for providing guidelines to download reference sequences from RefSeq in adequate formats in the ‘Opiniomics’ blog. The wild bird data set was developed with the contribution of Dr. A. C. Hurt and Dr. M. Klaassen. The WHO Collaborating Centre for Reference and Research on Influenza is funded by the Australian Department of Health. E.C.H. is funded by an Australian Laurate Fellowship (FL170100022). V.R.M. and T.C.S are Sydney Medical School Foundation Fellows. O.L and P.T.L.C.C. were supported by the European Union’s Horizon 2020 research and innovation programme under grant agreement no. 643476 (COMPARE), and The Novo Nordisk Foundation (NNF16OC0021856: Global Surveillance of Antimicrobial Resistance). This study was funded by an NHMRC Centre of Excellence grant # APP1102962.

