## Supplementary figure S1 for "CCMetagen: comprehensive and accurate identification of eukaryotes and prokaryotes in metagenomic data"

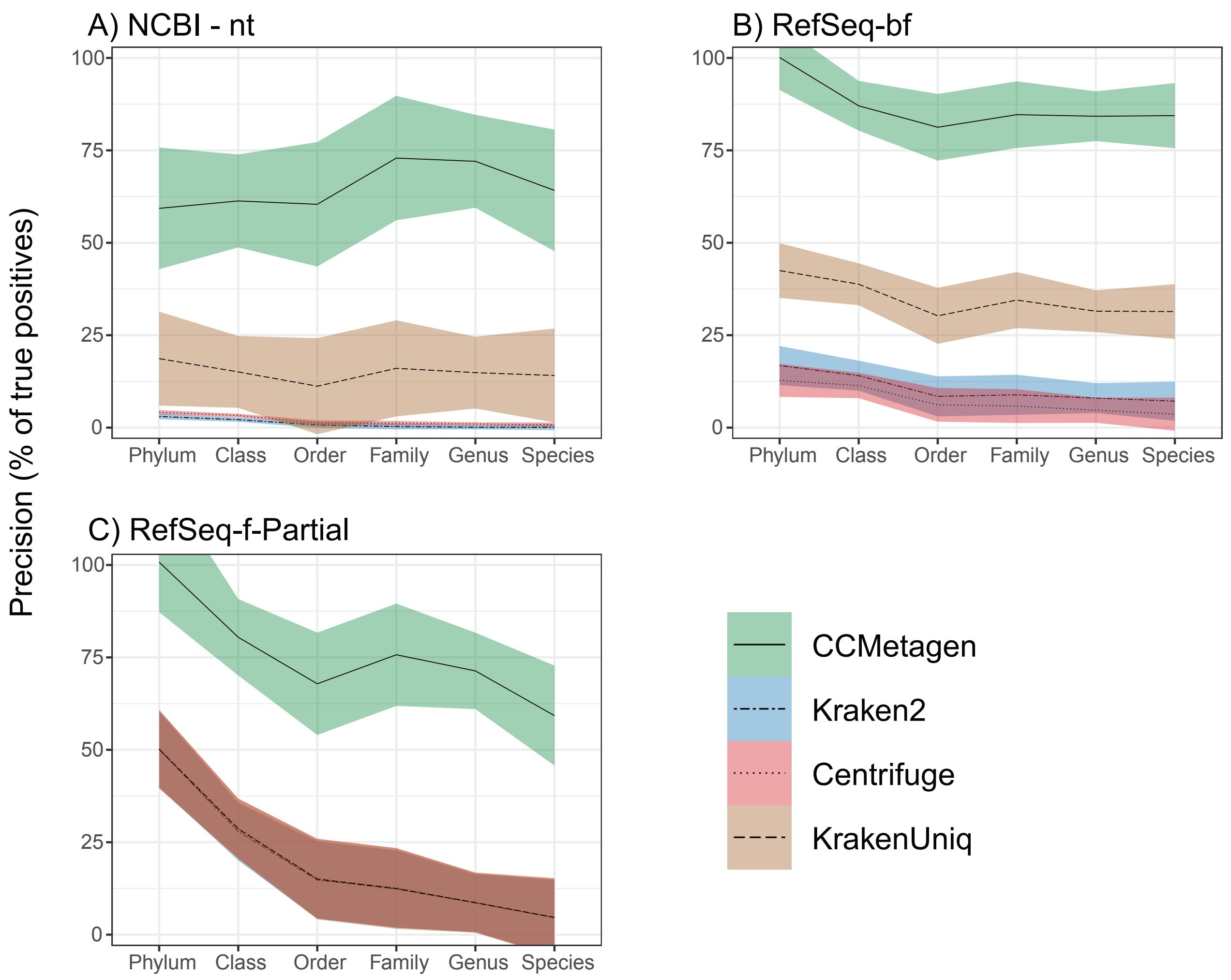


**Supplemental Figure S1.** Precision of the different methods, using three reference databases. (A) Results using the whole NCBI's nt collection as a reference database. (B) Results using the RefSeq (bacteria fungi) database, containing all bacterial and fungal genomes available.
(C) Partial RefSeq (fungi - partial) database, which mimics the effects of dealing with species without representatives in reference datasets. Kraken2, Centrifuge and KrakenUniq have overlapping results when using the Partial RefSeq database.
Shaded areas indicate 75% confidence intervals.
