## Supplementary figure S2 for "CCMetagen: comprehensive and accurate identification of eukaryotes and prokaryotes in metagenomic data"

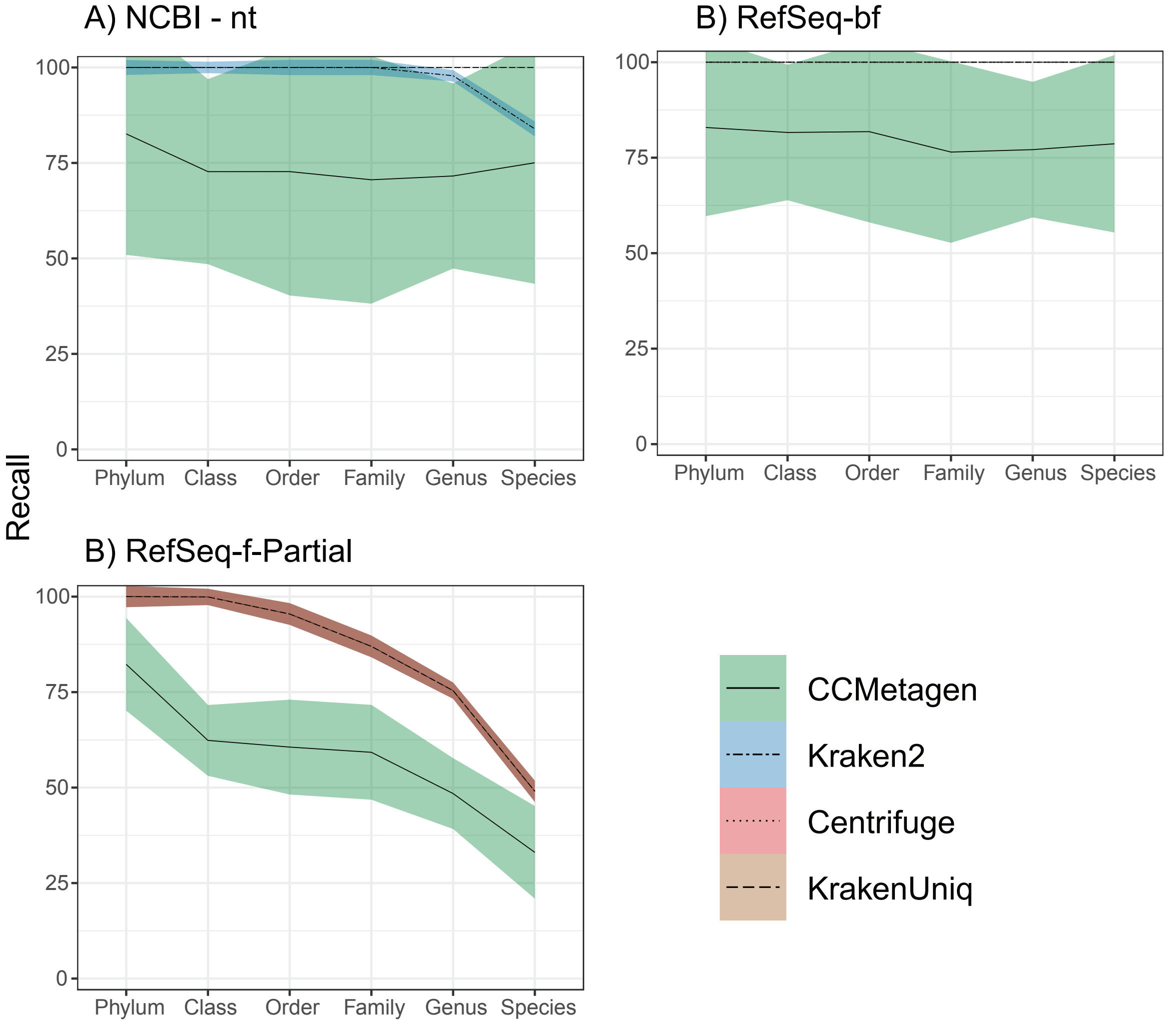


**Supplemental Figure S2.** Recall (% of taxa identified) of fungal taxa from a metagenome and a metatranscriptome test dataset. Note that the recall of Centrifuge, KrakenUniq and Kraken2 (up to genus level) is 100% (one straight line on top of graphs A and B).
The brown shaded area in C represents the overlapping results of Kraken2, Centrifuge and KrakenUniq. Shaded areas indicate 75% confidence intervals.
