## Supplementary material for "CCMetagen: comprehensive and accurate identification of eukaryotes and prokaryotes in metagenomic data": Sup Materials

**1. Reference databases:**

*1.1 RefSeq databases download:*

Fungal genomes (all assembly levels) and bacterial genomes (complete only) were downloaded from the NCBI website (<ftp://ftp.ncbi.nih.gov/blast/db/FASTA/nt.gz>).

We used the scripts described in <http://www.opiniomics.org/building-a-kraken-database-with-new-ftp-structure-and-no-gi-numbers/> to download sequences in a convenient format (*i.e.* containing taxids in

the sequence header) for Kraken2 and KMA (post-processed with CCMetagen).

For Centrifuge, we used their own download commands:

centrifuge-download -o library -a "Contig" -m -d "fungi" refseq > seqid2taxid.map

centrifuge-download -o library -a "Scaffold" -m -d "fungi" refseq >> seqid2taxid.map

centrifuge-download -o library -a "Chromosome" -m -d "fungi" refseq >> seqid2taxid.map

centrifuge-download -o library -m -d "fungi,bacteria" refseq >> seqid2taxid.map

centrifuge-download -o taxonomy taxonomy

cat library/*/*.fna > input-sequences.fna

To construct the RefSeq database containing only part of the fungal species analyzed (RefSeq-f-part), we manually removed sequences from 15 fungal species from the data sets (Supplemental table S5). This resulted in a database containing 15 of the 30 species in the fungal metagenome sample, and 7 of the 15 species in the metatranscriptome sample.

*1.2 NCBI nucleotide database download:*

Centrifuge: the official ncbi_nt non-redundant database for centrifuge, already indexed, was downloaded from the Centrifuge website.

Kraken2: we used the commands: kraken2-build --download-library nt --db nt

KMA/CCMetagen and KrakenUniq: we downloaded the partially non-redundant nucleotide database from the NCBI website (<ftp://ftp.ncbi.nih.gov/blast/db/FASTA/nt.gz>). This database was formatted to include taxids in sequence headers with custom scripts to be processed it with CCMetagen.

*1.3 Indexing databases:*

The databases were indexed as follows:

Centrifuge: centrifuge-build -p 10 --conversion-table seqid2taxid.map --taxonomy-tree taxonomy/nodes.dmp --name-table taxonomy/names.dmp input-sequences.fna ref_db_name

Kraken2 RefSeq databases: kraken2-build --build --db RefSeq_db --threads 10

Kraken2 nt database: kraken2-build --build --db nt --protein --threads 10

KrakenUniq:

krakenuniq-build --add-to-library input_sequences.fna --db ref_db_name

krakenuniq-build --db ref_db_name --threads 10 --kmer-len 31

KMA:

Indexes for KMA can be built on a Sparse mode, which limits the index to only include *k*-mers with a certain prefix, reducing processing time and memory consumption. On average, this will only save non-overlapping *k*-mers when a prefix of length 2 and *k*-mer size of 16 is chosen.

Note, however, that this step also reduces accuracy. Considering the size of the databases, we used all possible *k*-mers on both strands (option -Sparse -) for RefSeq, and only the ones prefixed with ‘TG’ for the NCBI nt collection. The databases indexed to function with KMA and CCMetagen are available at <https://cloudstor.aarnet.edu.au/plus/s/Mp8gLimDYoLfelH>, and were indexed as follows:

KMA RefSeq databases: kma_index -i refseq_bf_taxids.fna -o refseq_bf -NI -Sparse -

KMA nt database: kma_index -i nt_taxid.fas -o ncbi_nt -NI -Sparse TG

**2. Data analyses:**

*2.1 Quality control:*

Quality control was performed with prinseq-lite (Schmieder and Edwards 2011). As different programs were used to simulate the metagenome and metatranscriptome samples, the resulting sample files were in different formats: the fungal metatranscriptome and the bacterial metagenomes were in fasta, while the fungal metagenome was in fastq format.

The quality control of the fasta files was restricted to filtering out sequences with more than 5 ambiguous positions (Ns):

prinseq-lite -fasta sample1_R1 -fasta2 sample1_R2 -out_good sim_metatrans_good -out_bad sim_metatrans_bad -ns_max_n 5

While the quality control of the fastq files involved filtering sequences with more than 5 Ns and average quality <= 25 (*i.e.* keep sequences unless >/= 27 wrong bases were inserted in the simulated metagenome).

prinseq-lite -fastq sample1_R1 -fastq2 sample1_R2 -out_good sim_metagen_good -out_bad sim_metagen_bad -ns_max_n 5 -min_qual_mean 25 -out_format 3

*2.2 Metagenome classification:*

The analyses using Kraken2, Centrifuge and KrakenUniq were performed with default values:

kraken2 --output sample1_out.tsv --report $sample1_report.tsv --db ref_database --threads 14 --paired sample1_R1 sample1_R2

centrifuge -x ref_database -f -1 sample1_R1 -2 sample1_R2 -S sample1_out --report-file sample1_out_report.tsv -p 16

krakenuniq --report-file sample1.tsv --db ref_database --threads 16 --paired sample1_R1 sample1_R2

The KMA analyses were performed with settings recommended in the CCMetagen manual.

For a paired-end sample, using the nt database, the commands were:

kma -ipe sample1_R1 sample1_R2 -o sample1_out -t_db nt -t 4 -1t1 -mem_mode -and -apm f

For a single fasta file (bacterial metagenomes), also using the nt database, the commands were:

kma -i sample1 -o sample1_out -t_db nt -t 4 -1t1 -mem_mode -and -ca

CCMetagen was subsequently run as:

CCMetagen.py -i sample1_out.res -r nt -o sample1_CCMetagen_result
